# *MRtrix3*: A fast, flexible and open software framework for medical image processing and visualisation

**DOI:** 10.1101/551739

**Authors:** J-Donald Tournier, Robert Smith, David Raffelt, Rami Tabbara, Thijs Dhollander, Maximilian Pietsch, Daan Christiaens, Ben Jeurissen, Chun-Hung Yeh, Alan Connelly

## Abstract

*MRtrix3* is an open-source, cross-platform software package for medical image processing, analysis and visualization, with a particular emphasis on the investigation of the brain using diffusion MRI. It is implemented using a fast, modular and flexible general-purpose code framework for image data access and manipulation, enabling efficient development of new applications, whilst retaining high computational performance and a consistent command-line interface between applications. In this article, we provide a high-level overview of the features of the MRtrix3 framework and general-purpose image processing applications provided with the software.

## 1 Introduction

The use of medical imaging technologies such as Magnetic Resonance Imaging (MRI) has rapidly expanded over the last decades, sparking development of dedicated digital image analysis techniques tailored to these often large imaging datasets (3 and even higher dimensional) (Dhawan, 2011). These methods include artefact correction, anatomy segmentation, quantitative feature extraction, and spatial image registration which allows for comparison of features in subjects over time, between (groups of) subjects and across populations. Development of medical image analysis techniques is largely driven by the academic research community. Open software platforms have the potential to enable translation of novel method developments to, e.g., MRI hardware vendors, clinicians and the biomedical research community.

*MRtrix3* has been designed as an open-source, modular software platform for medical image analysis and visualization. This includes a lightweight framework for application development that facilitates efficient image access and manipulation, convenient multi-threading primitives, and support for a broad range of image file formats. End-user applications provided with *MRtrix3* include general-purpose image conversion and manipulation tools and the visualisation tool *MRView* tailored to multi-dimensional (3D+) medical image data. All components are designed with a strong focus on performance and memory efficiency, which are essential features for dealing with these often very large datasets. Computationally demanding quantitative imaging algorithms often require run times on the order of minutes (or hours), which may easily become days (or weeks) if performance is not optimised to deal with this specifically. In addition, *MRtrix3* offers support for external modules that can be built and distributed independently.

*MRtrix3* was historically developed with a particular emphasis on investigating brain white matter using diffusion-weighted MRI (dMRI), a medical imaging modality that can act as an indirect probe of neural microstructure, by measuring local hindrance of water diffusion (Johansen-Berg and Behrens, 2013; Jones, 2010). While it derives from the now deprecated MRtrix 0.2.x software package (Tournier et al., 2012), it has evolved massively over the years, both in terms of the underlying codebase, and the scope of the functionality provided. The platform hence incorporates a number of features tailored to the orientational nature of dMRI data within each imaging voxel, such as a representation using spherical harmonics basis functions, and a bespoke file format for 3D streamlines. *MRtrix3* also provides a wide range of dMRI analysis methods, outlined in the *Diffusion analyses* section below. Thanks to these features, *MRtrix3* has primarily gained support in the dMRI community. However, its architecture facilitates image processing in general, and *MRtrix3* also offers image analysis methods less specific to dMRI such as image filtering and calculation, statistics, image denoising (Veraart et al., 2016) and Gibbs ringing suppression (Kellner et al., 2016), intensity normalisation (Raffelt et al., 2017a) and diffeomorphic image registration (Raffelt et al., 2011).

In this paper, we describe the guiding principles underlying the design and implementation of *MRtrix3*. We outline the main aspects of the software architecture, particularly the image access and multi-threading primitives, as well as the build system and related developer tools. Finally, we showcase the visualisation capabilities in *MRView* and give a brief overview of the currently supported tools for general-purpose image processing and dMRI analysis.

## 2 Guiding principles

### 2.1 Reproducible neuroscience

With a growing number of neuroimaging studies and the increasing prevalence of advanced data analysis in radiology practice, reproducibility becomes ever more important. Publishing open and reproducible analysis pipelines enables neuroscientific research to replicate pilot results on larger datasets, and also facilitates comparing sensitivity and specificity of various methods. In addition, publicly available software can help the clinical translation of novel analysis techniques.

The *MRtrix3* collective believes that reproducible neuroscience relies first and foremost on open source software. Publishing source code, tracking versions, and documenting improvements and bug fixes, is vital to ensure full transparency on what was done in a study. We are therefore fully committed to keeping *MRtrix3* open source and tracking the version history using Git. The software is published under the terms of the open source Mozilla Public Licence 2.0^1^, which permits third-party modification, distribution and commercial use as long as the source code is disclosed. Furthermore, *MRtrix3* also incorporates continuous integration testing to detect and avoid adverse effects of code changes between versions.

In addition, *MRtrix3* provides the requisite functionalities for handling data in the Brain Imaging Data Structure (BIDS)^2^, a recent standard for organising and sharing neuroimaging data (Gorgolewski et al., 2016). These include importing and exporting dMRI gradient encoding and other metadata in JSON and BIDS bval/bvec format, ultimately enabling integrating *MRtrix3* commands in BIDS Apps (Gorgolewski et al., 2017). Finally, *MRtrix3* also offers automatic history tracking in the output file header, archiving the pipeline of operations used to generate each output image, as well as the exact version tag of each invoked command.

### 2.2 Documentation & support

User documentation is available in the inline help page of each command and also on a dedicated documentation website^3^ that includes additional background information, tutorials and pipelines. The documentation is kept up to date with the current release and can be retrieved for older versions. Major updates and new software releases are also announced on the main website and blog^4^.

Another key benefit to open development is the interaction between users and developers as one community. To foster these interactions *MRtrix3* provides an online forum^5^, where a growing number of users post questions, receive support, and have scientific discussions. Furthermore, users and developers can report and track technical issues through Github^6^, and suggest new features for future releases. Github is also the platform where both internal and external developer contributions are discussed and managed.

## 3 Design aspects

*MRtrix3* has been designed from the outset to facilitate the development of high-performance applications concerned with the processing and analysis of medical image datasets, and to offer a consistent and simple command-line interface. It focuses primarily on command-line applications to provide a wide range of functionalities that can be combined and scripted to form complex automated workflows. It relies on modern standards and technologies, particularly C++11^7^ and *OpenGL^8^* 3.3, yet strives to limit the number of external dependencies to a small and well-supported set, available across a broad range of platforms; currently these include Python^9^ (for building the software and using scripts), *zlib^10^* (for access to compressed images), *Eigen3^11^* (for high-performance linear algebra support), and *Qt^12^* (version 4 or 5, for graphical user interface support). All of these dependencies are available across a wide range of platforms, allowing *MRtrix3* to run natively on most flavours of Unix, including GNU/Linux and macOS as well as a range of high-performance computing (HPC) environments, and Microsoft Windows via the *MSYS2^13^* project or Windows Subsystem for Linux (*WSL*)^14^.

Most of the functionality is written using C++ templates, providing high runtime performance by allowing the compiler to optimise away any redundant overhead. Motivated by a heavy focus on multidimensional image access, simple yet powerful constructs for multi-threaded operations are provided. The framework provides native access to and seamless interaction between a range of image file formats commonly used in the medical imaging community, including native support for DICOM^15^ (read-only), Analyze^16^, NIfTI1^17^, NIfTI2^18^, MGH^19^, and *MRtrix3*’s own dedicated image formats^20^, as well as compressed versions thereof.

While most *MRtrix3* commands consist of compiled binary executables, some are written in Python, and typically invoke other *MRtrix3* executables to perform more complex tasks. Moreover, a number of scripts invoke commands from 3rd party software packages such as FSL (Jenkinson et al., 2012) or ANTs (Avants et al., 2009), providing convenience wrappers to perform specialised tasks (see the *Image Processing tools and analyses* section below for details).

### 3.1 Image access

Functionality for image input/output is provided by the core *MRtrix3* library, so that all *MRtrix3* applications can read and write all supported image formats directly: there is no requirement to convert the data to and from specific image formats. All *MRtrix3* applications consistently use the same coordinate system, which is identical to the NIfTI standard. *MRtrix3* handles coordinates consistently across file formats and in addition, the native *MRtrix3* image format allows automated handling and storage of coordinate system dependent information such as the dMRI gradient table in the image header to ensure this information remains consistent throughout the analysis pipeline.

#### 3.1.1 Optimised data access

Access to image data is automatically optimised using a variety of techniques to ensure near-optimal performance across a range of operating conditions. The various strategies employed are outlined in this section.

By default, image access is achieved via memory-mapping with on-the-fly data type conversion. However, when the stored data type matches the data type requested by the application, *MRtrix3* commands will automatically use “direct IO” access, where data are accessed as a native array in memory, thus avoiding the overhead of the function calls otherwise necessary for the conversion. When desired, applications can also explicitly request “direct IO” access to the data: this is particularly beneficial for applications that will access image data repeatedly, for example, multi-pass algorithms, patch-based operations, interpolation, etc. To achieve this, the data are loaded directly into RAM in the desired data type, unless they are already stored as such (in which case memory-mapping already provides “direct IO” access). Furthermore, while image writing is also typically done using memory mapping, there are cases where doing so incurs a loss of performance. This is particularly relevant when operating on networked file systems where random writes to memory-mapped regions can result in heavy network usage. In this case, *MRtrix3* uses an alternative *delayed write-back* strategy, with the backend holding the data in RAM and automatically committing the data to storage when the image is closed.

*MRtrix3* also has a flexible concept of *strides*, allowing the multi-dimensional image data to be stored in RAM (or file when the image format supports it) in any reasonable order (see Appendix A for details). It is for example entirely possible to have the data stored in a volume-contiguous manner (for 4D files), meaning that the data for all volumes of a given imaging voxel are co-located in RAM (see Fig. A1). This allows for efficient access and maximises CPU cache reuse, in some cases improving performance considerably. Developers can specify these strides when accessing images with a particular purpose in mind (e.g., voxel-wise processing of 4D files), and the *MRtrix3* backend will load and re-order the image data in RAM if necessary, but still use “direct IO” access via memory-mapping where possible to avoid needless data copies. Images can be stored in any stride ordering supported by the chosen file format. Of the supported formats, only *MRtrix3*’s own formats support arbitrary stride ordering, a feature that offers distinct performance advantages when processing different types of data, particularly by allowing volume-contiguous storage.

#### 3.1.2 DICOM handling

*MRtrix3* includes its own fast DICOM import handling, allowing all *MRtrix3* applications to seamlessly support DICOM images as input. The DICOM import code handles data stored using the recent DICOM multi-frame standard, as well as the standard multi-file storage, including Siemens mosaics. If present, it will also extract the diffusion MRI encoding information for 3 major vendors (Siemens, GE, Philips), and make it available to the application as part of the image header. Other types of information are also extracted if found, including the EPI phase-encode direction and slice timings.

DICOM images can be provided as a single .dcm file, or more commonly, as a folder containing the multiple DICOM files that make up the full DICOM series. The import code will scan through all files in the folder, recursing into any subfolders encountered, and collate all the information into a hierarchy by patient / study / series / images. If the folder contains a single DICOM series, the data are loaded as a single dataset directly with no further interaction required. Otherwise, the user will be presented with a listing of the contents and required to select the desired series.

### 3.2 Performance and piping

#### 3.2.1 Unix pipes

*MRtrix3* allows temporary images to be created in RAM for use as scratch buffers within applications and also passed between applications via Unix pipes. This allows the software to be used in flexible, powerful combinations, for example:

~~~
$ mrthreshold input.mif -abs 0.5 - | maskfilter - erode - | mrview input.mif -overlay.load -
~~~

Listing 1: illustration of the use of Unix pipes in MRtrix. The command above will threshold image “input.mif” at an absolute value of 0.5; erode the resulting binary mask by one voxel; visualise the original input image, with the mask calculated in the previous step overlaid on top

Rather than passing the full image data through the pipe, temporary images are created to hold the data, and only the filename is passed through the pipe. This helps to ensure optimal performance since the data can be accessed via memory-mapping at both ends of the pipe, eliminating any needless data copies, and keeping RAM usage to a minimum.

#### 3.2.2 Multi-threading

Multi-threading is a type of computer execution technique supported by modern hardware that allows a program to run across multiple threads of execution concurrently. Each thread executes independently while sharing its resources and coordinating with the other threads in the process. In *MRtrix3*, almost all commands have been multi-threaded, by default running with the maximum utilization of CPU processors to achieve high computing performance. In addition, *MRtrix3* provides a powerful, yet simple to use framework for developers to write multi-threaded operations, particularly for iterating over all voxels in an image, or for multi-stage pipelines.

Multi-threaded looping is demonstrated in the example in Appendix B, illustrating the simplicity of use. In this implementation, concurrent threads will typically process adjacent rows of data; this is done to ensure optimal performance by maximising cache reuse, minimising disk seek latency, and maintaining a sufficiently fine level of granularity to avoid idle CPU cores. By default, the order of traversal of the image axes is set by the strides of the image, although this can be specified explicitly by the developer if desired. Moreover, the developer can control which images are to be looped over simultaneously, which axes to iterate over, and of these, which are to be processed within each thread (a single row by default).

### 3.3 Developer tools

*MRtrix3* is developed using modern software practices with transparency in mind. Software version control is managed using Git, hosted openly on Github^21^. Issues are tracked and discussed publicly on GitHub^22^. Most *MRtrix3* commands have tests to ensure the output remains consistent with expectation throughout the development cycle. Test data are hosted on a separate Git repository^23^ to keep the main source code repository small and lightweight. Continuous integration testing is managed by TravisCI^24^ (for Linux and macOS) and AppVeyor^25^ (for Microsoft Windows).

*MRtrix3* is designed to minimise the amount of effort required to develop complete image processing applications, as illustrated in Figure 1. Developers have access to a range of classes for seamless access to images and other types of data, facilities for multi-threading operations over images and for parallel processing of streams of data, image filters and other convenience methods. The core API documentation is available online^26^, and is generated directly from the code using Doxygen^27^.

**Figure 1:**
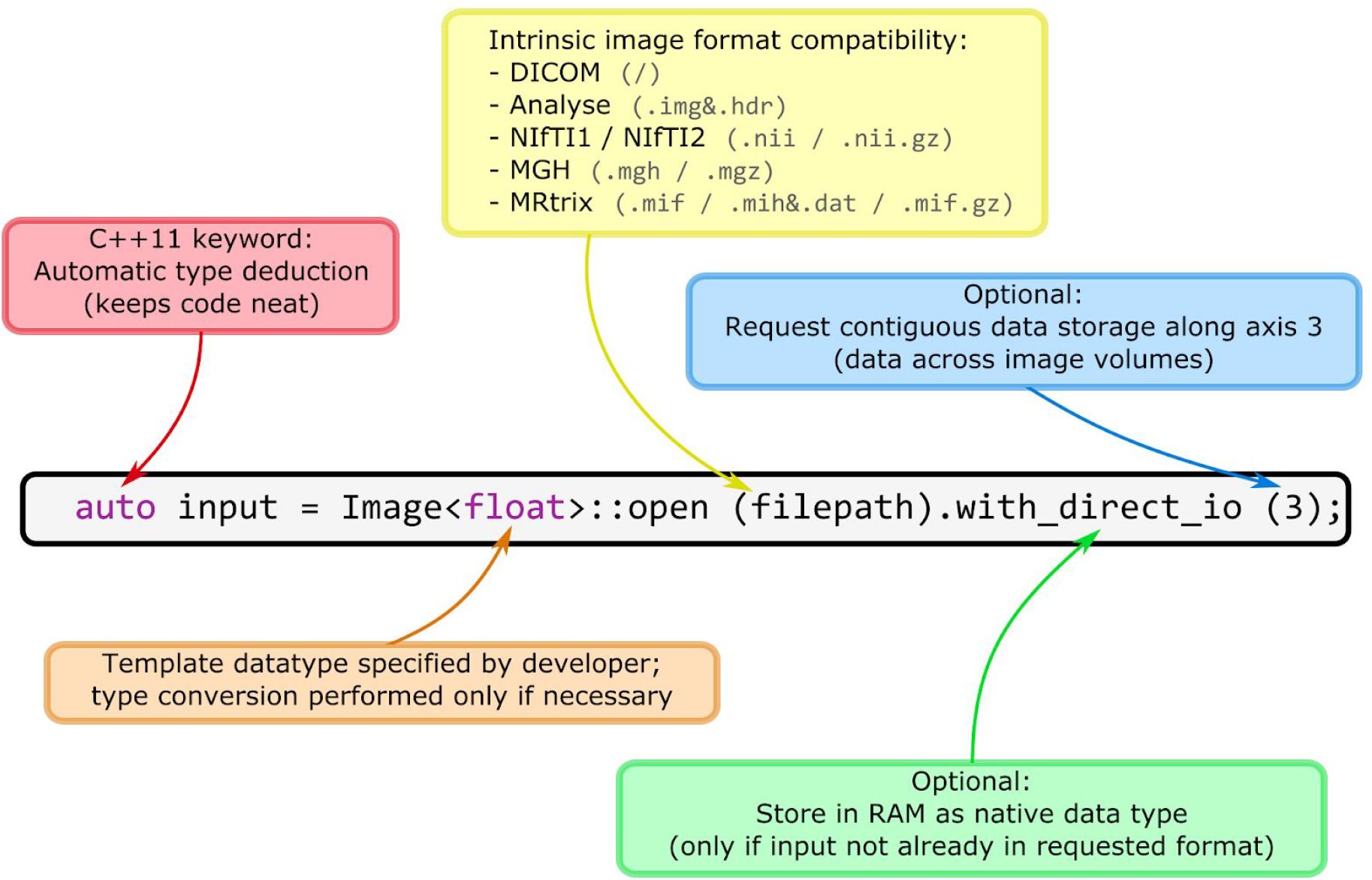
Demonstration of features provided within a single line of code when using the *MRtrix3* software library to access an image from the filesystem.

The development tools and build system are also demonstrated in an example application in Appendix B (other examples are available in the developer documentation^28^).

#### 3.3.1 Integrated command-line interface and documentation

*MRtrix3* provides a framework for simultaneously documenting, specifying and retrieving command-line arguments and options, all centralised within a single function for each application. This includes authorship and copyright information, a general description of the command and specific descriptions of the arguments and options, and any relevant external references. This information is used to generate the application’s online help page, and at runtime to display its help page, to automatically check the validity of the arguments, and to retrieve these arguments and options at any point in the code. See the code example in Appendix B for an illustration of this framework.

#### 3.3.2 Build system

The software includes a bespoke system to manage configuring and building the software. It is comprised of two Python scripts: configure and build. The former detects various aspects of the operating environment (OS, availability of C++11 compliant compiler and required libraries, etc). The build script scans the code to build a full dependency tree based on simple rules about file location and naming conventions, identifies outdated targets, assembles a list of tasks required to bring the targets up to date, and then launches the relevant commands in the right order and in parallel for fast build times. This allows the code to be modified without requiring any other changes (for example, to external Makefiles): a new command can simply be dropped into the right folder, and its corresponding binary will be compiled the next time build is invoked.

#### 3.3.3 External contributions

*MRtrix3* encourages users to contribute their own methods to the software and recognises their need to distribute these however they see fit. Therefore, the *MRtrix3* build system provides a convenient primitive for external modules that enables developers to build their own applications, linked to the core *MRtrix3* library and leveraging its image access and multi-threading primitives. These external modules can be distributed separately under any open source license or may be integrated into a future *MRtrix3* release. In either case, developers are given full credit for their work through embedding of author information, external references, and copyright statement, within the code and resulting command help page and documentation.

#### 3.3.4 Python scripts

While piping of images (as described in Unix pipes (section 3.2.1)) can be used to chain several *MRtrix3* commands into composite applications, *MRtrix3* also contains a dedicated Python scripting library to aid the development of more complex applications and processing pipelines involving multiple *MRtrix3* (or other) binaries. Features of the scripting library include standard command-line options, help page formatting and documentation generation consistent with *MRtrix3* binary commands, scratch directory management, and the ability to continue from any point in a previous execution for long or computationally demanding pipelines. The application module and library functions on which these scripts are built are additionally designed to be both accessible and functional for researchers looking to construct their own processing pipelines from existing commands and to interface with external tools.

## 4 MRView

The *MRtrix3* viewer, *MRView*, was written to be cross-platform, high performance, convenient to use, and extensible. To meet these aims, *MRView* uses the high-performance industry-standard OpenGL^29^ 3.3 API to leverage the graphics capabilities of modern systems, and the widely supported and open-source Qt toolkit^30^ to manage the interface elements (widgets). It is also designed from the outset for interactive use from the command-line. This allows users to rapidly experiment with new ideas, develop and debug pipelines, and inspect the output of processing commands. Two aspects of its design play a role in supporting this: first, *MRView* accepts input from Unix pipes (as illustrated in the example in section 3.2.1); and second, it is quick to launch and display (through a number of design decisions, including the use of memory-mapping for “lazy loading” and delayed initialisation of interface elements until the point of use). See the *MRtrix3* website for demonstration videos^31^.

An important aspect of *MRView* is its ability to load multiple images concurrently, allowing fast switching between them via the menu or the PageUp / PageDown keys. In combination with the optimisations mentioned above, this allows interactive use in otherwise demanding workflows, such as loading a long list of images into the viewer concurrently and inspecting them with interactive framerates, even when the data are held on network file shares. This type of action is useful to quickly scan through the results of a processing step over all participants in a large study (to check the quality of the image masks, for example).

*MRView* is also capable of displaying slice-wise information at arbitrary oblique cuts through the data. This is particularly useful to display features such as streamlines obtained via tractography in their most appropriate anatomical context, for instance. Off-axis rendering is available in all viewing modes (see below). *MRView* can also handle higher-dimensional images, currently up to 5D. Navigating between volumes (4th dimension) or volume groups (5th dimension) is simple and fast via the arrow keys, the menu, or the View tool (see below).

A core principle in *MRView* is that content is always displayed at the correct location in *scanner* coordinates, not relative to the *image* voxel coordinates. This applies to the main image, but also overlays and streamlines, and any directional information such as vectors or orientation density functions (ODFs). This allows the concurrent display of different types of data, independently of how these data were generated. For instance, images can be overlaid together even if their voxel sizes and/or orientations differ, streamlines produced by tractography can be displayed over other images than those used to produce them (e.g. co-registered anatomical images), ODFs or vectors will point in the expected direction irrespective of the image currently displayed, etc.

*MRView* provides a number of different viewing ***modes***, including single-slice; orthoview (a montage of 3 orthogonal slices), lightbox (a grid montage of adjacent slices or volumes); and volume render (3D rendering using ray-tracing). Figure 2 shows a screen capture of *MRView* in volume render mode.

**Figure 2:**
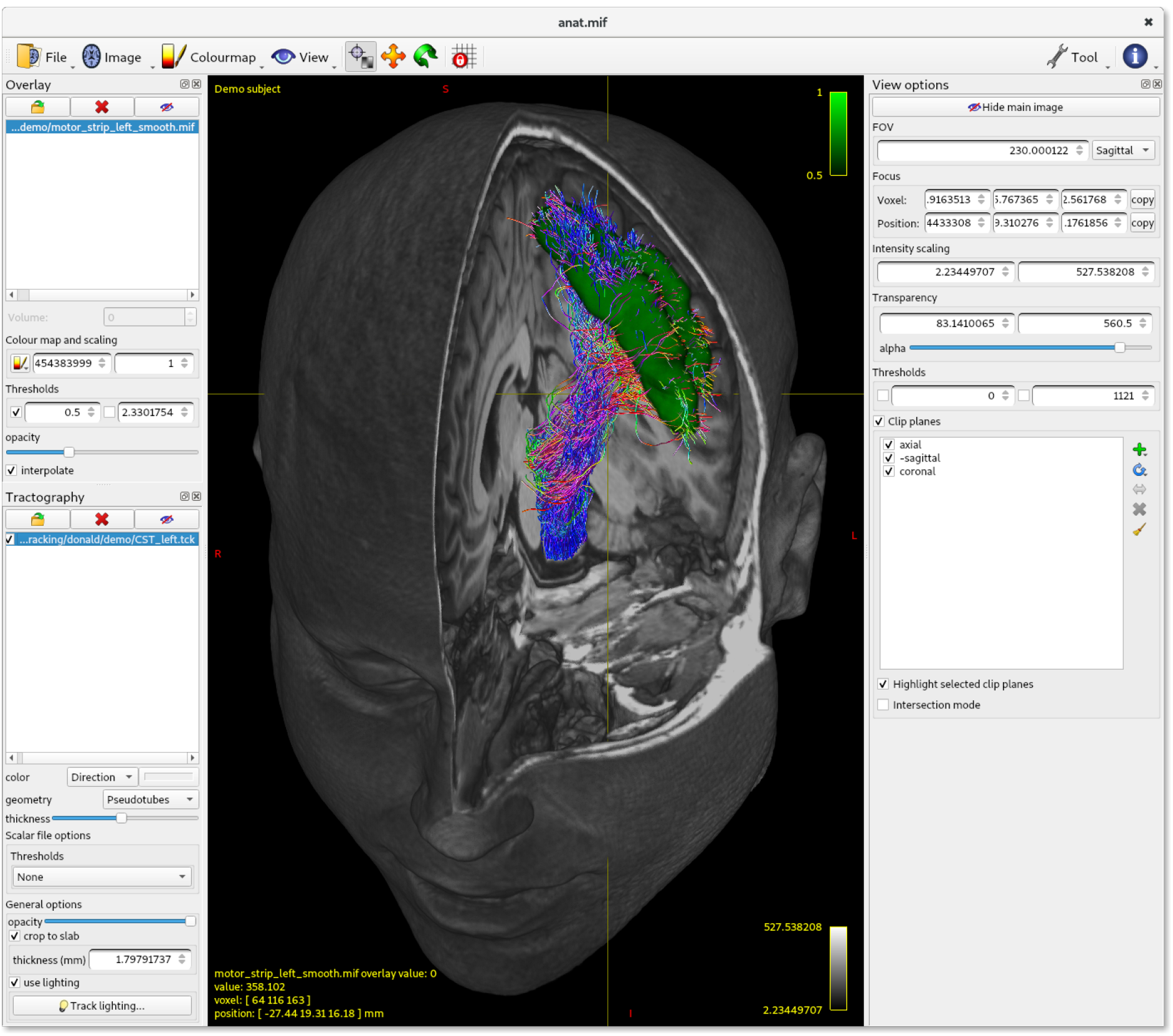
MRView in volume render mode, showing an anatomical T1-weighted MRI with 3 clip planes, with a region of interest corresponding to the motor strip rendered using the overlay tool (in green), and probabilistic streamlines delineating the corticospinal tract shown via the tractography tool.

These modes are supplemented by a number of independent ***tools*** providing additional specific functionality. These include:

- **View**: provides fine control over generic aspects of the display, such as brightness/contrast, location of crosshairs, and access to mode-specific settings such as clip planes for the volume render.
- **Overlay**: overlay additional images on top of the main image, with control over colour map, transparency, and thresholds.
- **ROI editor**: provides an interface to edit binary mask images.
- **Tractography**: visualisation of tractography output, with options for streamtube rendering.
- **ODF render** tool: visualisation of orientation density functions (Figure 3), provided as spherical harmonic coefficients, samples along a set of fixed orientations (‘dixels’^32^), or tensor coefficients.
- **Fixel plot** tool: display voxel-wise vectors, provided either as 4D images of X,Y,Z coefficients or using *MRtrix3*’s sparse fixel storage format^33^ (Figure 4). See (Raffelt et al., 2017b) for a description of the fixel concept.
- **Connectome** tool: visualise nodes and edges obtained through connectome analysis, using a variety of techniques (Figure 5).
- **Screen capture** tool: write the contents of the main display to file using the widely supported open-source Portable Network Graphics (PNG) format^34^. This additionally allows the creation of movies (see for example demonstration videos on the *MRtrix3* website, and the supplementary materials provided with (Mito et al., 2018)).

**Figure 3:**
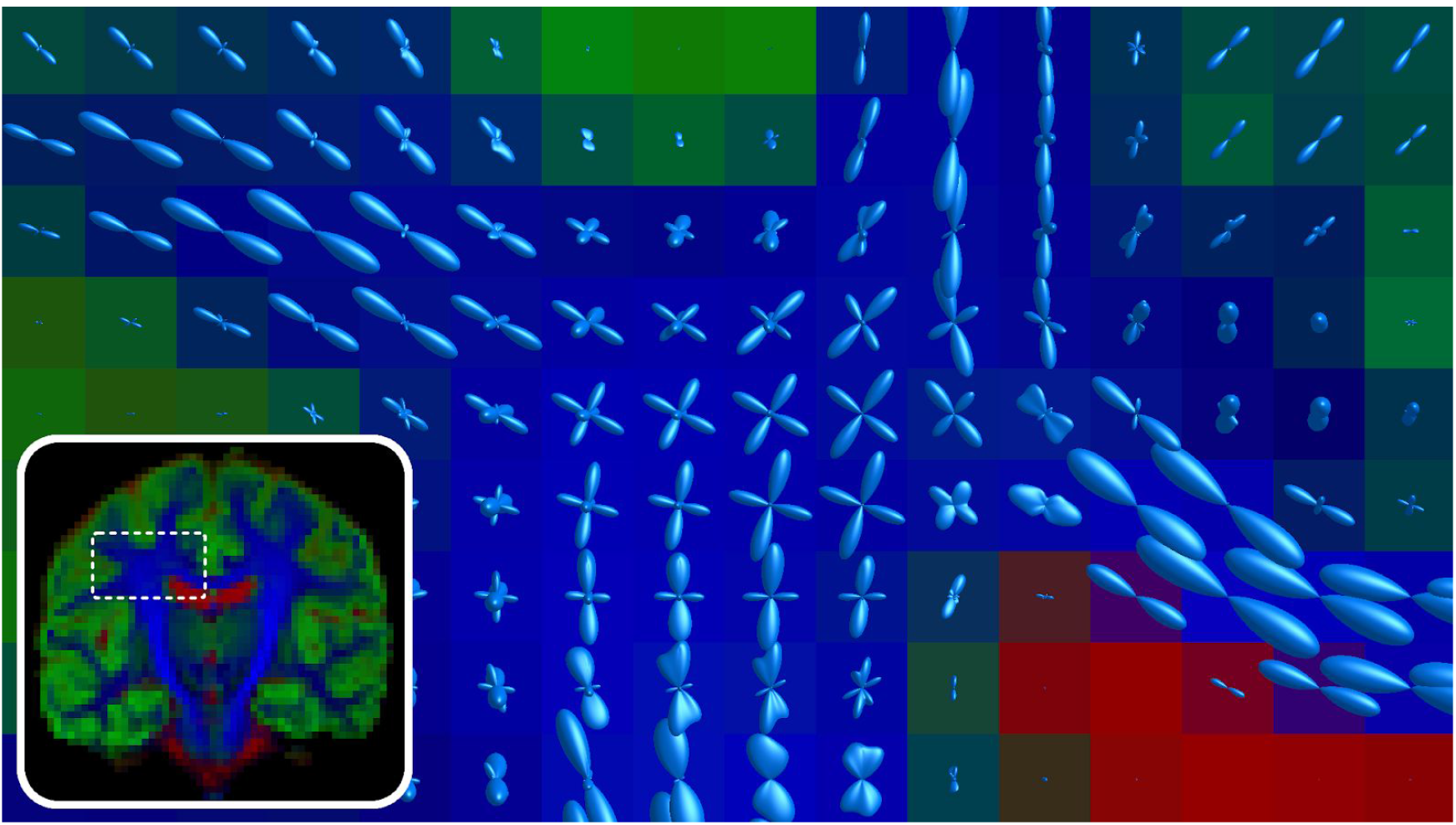
Multi-tissue Constrained Spherical Deconvolution result using 3 tissue types: WM-(blue), GM-(green) and CSF-like (red) compartments, with WM fibre orientation distributions (lighter blue) overlaid.

**Figure 4:**
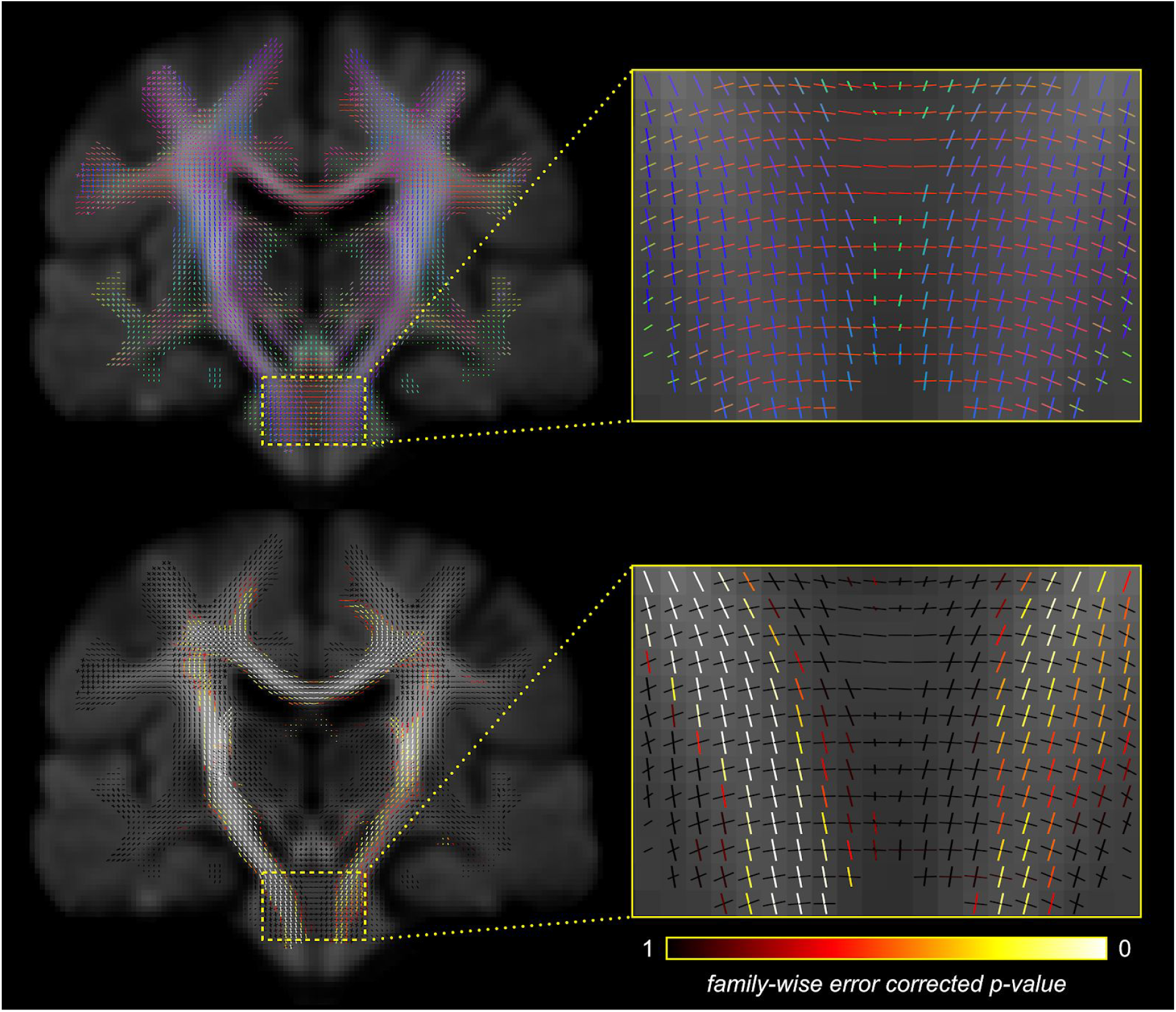
An example of fixel-based analysis results displayed using *MRView*’s Fixel plot tool. This example shows the comparisons of fixels between motor neuron disease patients and healthy controls, with the use of connectivity-based fixel enhancement. Top row: Fixels are presented in directionally-encoded colours, red: left-right, green: anterior-posterior, blue: inferior-superior. Bottom row: Fixels are coloured based on family-wise error corrected *p*-value, highlighting the fixels with a significant fibre density loss in the patient group. This figure is adapted from (Raffelt et al., 2015) with copyright permission.

**Figure 5:**
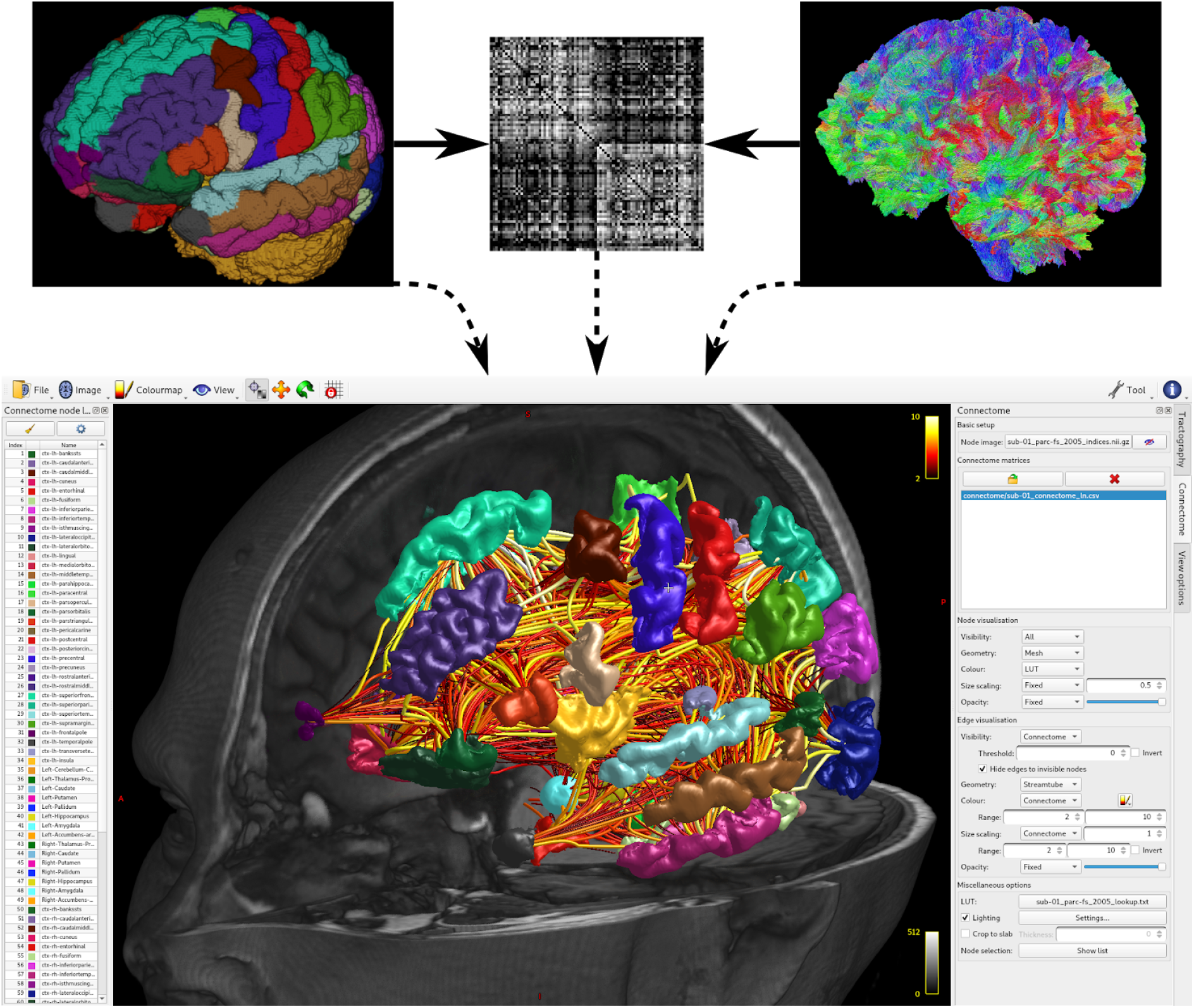
Generation of structural connectome matrix, and subsequent visualisation using the *MRView* Connectome tool. A brain parcellation from any of a range of data/software sources can be combined with a whole-brain tractogram generated using *MRtrix3*’s advanced tractography methods to produce a connectome matrix encapsulating a measure of connectivity between every pair of brain regions; the *MRView* connectome tool provides a range of features for displaying and navigating connectome data and/or the results of network-based statistical inference.

## 5 Image processing tools and analyses

*MRtrix3* comes bundled with a range of end-user applications for performing general-purpose image manipulations as well as complex analyses, with a strong (although not exclusive) focus on diffusion MRI. *MRtrix3* command names follow a convention to reflect the purpose of the command and aid discoverability. Firstly, a set of shorthand “codes” are defined for the main types of data. These indicate directly within the command name itself the type of data operated on by the command; for example, “`mr`” for generic (magnetic resonance) images, “`dwi`” for diffusion-weighted images and “`tck`” for tracks.

Using these, two standardised formats of command names exist:

- “`DataOperation`”, where the type of data input to the command and the operation to perform are concatenated; for example, “`mrconvert`” for converting images to different formats, or “`mrinfo`” for providing information about the image header.
- “`Data2Data`”, where the command involves some form of conversion from one type of data to another; for example, “`warp2metric`” for computing different metrics from spatial warps (deformation fields).

### 5.1 Basic image operations

*MRtrix3* provides a suite of commands for performing various common manipulations of image data. Each command is small, yet powerful and modular. This allows for performing a wide range of common processing tasks whilst providing maximal flexibility to the user. These basic commands include: querying image header information (`mrinfo`); converting between image formats, including data exclusion / axis permutation / header manipulation (`mrconvert`); concatenating images along any dimension (`mrcat`); a voxel-wise image calculator, allowing for simple as well as complex mathematical expressions (`mrcalc`); computing summary statistics (e.g. mean, std. dev., etc.) across different images or along image axes (`mrmath`); and generating summary statistics of image intensities within regions of interest (`mrstats`);

### 5.2 Diffusion analyses

*MRtrix3* includes tools to undertake different types of diffusion analyses from start to finish. In particular, it includes state-of-the-art tools for pre-processing, voxel-level modelling, fibre tracking and connectomics, and groupwise analysis. The main technologies currently included in *MRtrix3* are listed below.

**Pre-processing tools:** MP-PCA denoising (Veraart et al., 2016); removal of Gibbs ringing artefacts (Kellner et al., 2016); a convenience wrapper for FSL tools to perform motion, eddy-current, and susceptibility-induced distortion correction (Andersson et al., 2017, 2003; Andersson and Sotiropoulos, 2016); and a convenience script to perform bias field correction, based on either ANTs N4 (Tustison et al., 2010) or FSL (Smith et al., 2004; Zhang et al., 2001).

**Voxel-level modelling:** diffusion tensor imaging (DTI) (Basser et al., 1994; Veraart et al., 2013); constrained spherical deconvolution (CSD) for single-shell fibre ODF estimation (Tournier et al., 2007, 2004) and multi-shell multi-tissue CSD for multi-shell data (Jeurissen et al., 2014), supported by several response function estimation algorithms (Dhollander et al., 2016; Tax et al., 2014; Tournier et al., 2013) (Figure 3).

**Tractography and connectomics:** deterministic tractography using DTI (Mori et al., 1999) or fODFs (Tournier et al., 2012); probabilistic tractography using DTI via the wild bootstrap (Jones, 2008) or fODF sampling (Jeurissen et al., 2011; Tournier et al., 2012, 2010); multi-shell multi-tissue global tractography (Christiaens et al., 2015); anatomically-constrained tractography (Smith et al., 2012); spherical-deconvolution informed filtering of tractograms (SIFT) (Smith et al., 2013) and its newer SIFT2 variant (Smith et al., 2015); tools to support connectomic analyses, including network-based statistics (NBS) (Zalesky et al., 2010) and its threshold-free variant (Vinokur et al., n.d.).

**Group-wise image analysis:** diffeomorphic registration of ODF images and construction of population templates (Raffelt et al., 2011); voxel-based analysis using permutation testing (Anderson and Robinson, 2001; Winkler et al., 2014) and threshold-free cluster enhancement (Smith and Nichols, 2009); and fixel based analysis (Raffelt et al., 2017b) using connectivity-based fixel enhancement (Raffelt et al., 2015).

**Visualisation and maps:** directionally-encoded colour (DEC) maps for tensors (Pajevic and Pierpaoli, 1999) and fODFs (Dhollander et al., 2015b); panchromatic sharpening (Dhollander et al., 2015a) and luminance correction (Dhollander et al., 2018); track density imaging (TDI) (Calamante et al., 2010), track-weighted imaging (TWI) (Calamante, 2017) and track orientation density imaging (TODI) (Dhollander et al., 2014).

## 6 Conclusion

*MRtrix3* was designed from the outset as a framework for the development and dissemination of high-performance imaging research applications, guided by the principles of open and reproducible science. While its historical focus has been on diffusion MRI analysis, the bulk of the functionality is designed to support the broadest range of applications possible. We encourage researchers and developers alike to consider using *MRtrix3* in their research, whether focussed on diffusion MRI or other imaging modalities.

## Acknowledgements

This work was primarily supported by the National Health and Medical Research Council (NHMRC) of Australia (Program Grants 400121, 628952, 1091593; AC research fellowship 1026077). We are additionally grateful to the Victorian Government’s Operational Infrastructure Support Program for their support.

This work also received funding from the European Research Council under the European Union’s Seventh Framework Programme (FP7/2007-2013/ERC grant agreement no. [319456] dHCP project), and was supported by the Wellcome/EPSRC Centre for Medical Engineering at King’s College London [WT 203148/Z/16/Z]; the Medical Research Council [MR/K006355/1] and by the National Institute for Health Research (NIHR) Biomedical Research Centre based at Guy’s and St Thomas’ NHS Foundation Trust and King’s College London. The views expressed are those of the author(s) and not necessarily those of the NHS, the NIHR or the Department of Health.

RS is recently supported by fellowship funding from the National Imaging Facility (NIF), an Australian Government National Collaborative Research Infrastructure Strategy (NCRIS) capability.

BJ is a postdoctoral fellow of the Research Foundation Flanders (FWO Vlaanderen), Grant no. 12M3116N & 12M3119N.

## 8 Appendix A: Image strides

Image data can be stored on file or in RAM in a number of ways. At heart, multi-dimensional image data consists of one value for each position, indexed by coordinates *x*, *y*, *z*, …. The first 3 coordinates conventionally refer to the voxel indices along the spatial *x*, *y*, *z* axes, and higher indices would refer to separate volumes, or any reasonable indexing thereof. However, when stored on file or in RAM, the data will be arranged linearly, and indexed by a single *offset*. A strategy is needed to manage access to the data by mapping voxel coordinates to file (or memory) offsets.

To illustrate the issue, consider the problem of storing the elements of a dense *m*×*n* matrix. In many linear algebra packages, matrices are stored in so-called *column-major* format: each element of the first column is located directly after the other until the end of that column, followed by the elements of the next column, and so on. Assuming this storage convention, elements that are adjacent along the column direction are stored at adjacent offsets in RAM (or file), while elements adjacent along the row direction are placed at least *m* elements apart. Hence, the element at (*i*,*j*) is at offset = *i* + *jm* relative to the element at index zero. This corresponds to *strides* of [1 *m*]: the step to find the next element along the column direction requires a stride of 1 element, while the equivalent step along the row direction requires a longer stride of *m* elements.

**Figure A1:**
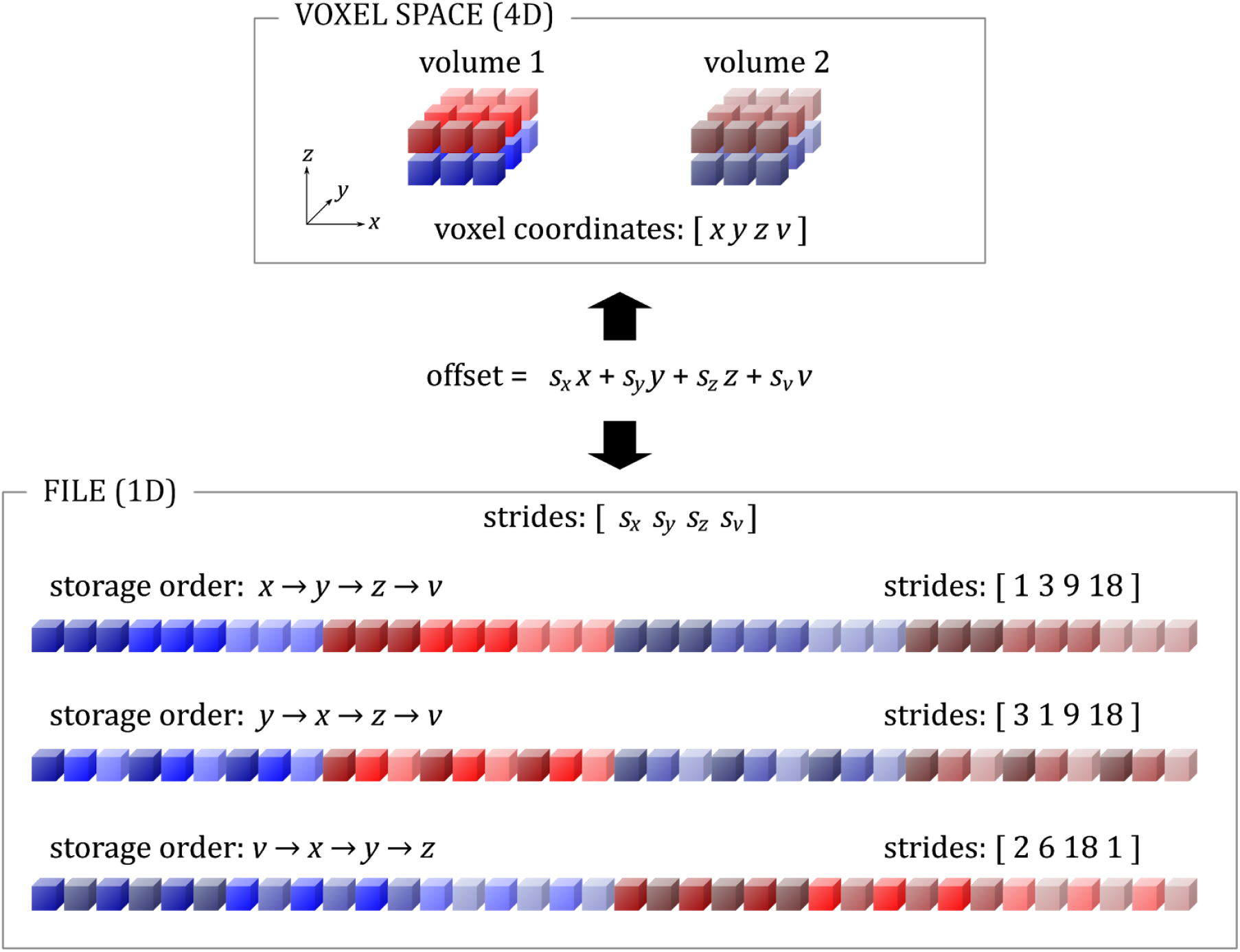
Illustration of the concept of strides when storing and accessing image data. The issue is to relate voxel coordinates [*x y z v*] (top box) to a single *offset* on file (bottom box). This can be done in a number of different ways. In the first case, data are stored in order of traversal along the *x* axis first, then each row along the *y* axis, then each slice along the *z* axis, and finally each volume. In the second case, the order is swapped between the *x* & *y* axes. The last case illustrates the case of volume-contiguous storage, where the data are traversed along the volume axis first, followed by the spatial axes.

However, matrices can be (and often are) also stored in so-called *row-major* format, where elements along the first row are stored contiguously, followed by the next row, and so on. In this case, the element at (*i*,*j*) is at offset = *in* + *j*; this corresponds to strides of [*n* 1]. Note that while two matrices stored in row-major and column-major format respectively may be logically equivalent, the *order* in which their elements are stored differs, and there is no reason to prefer one convention over the other. In fact, many implementations support both, since it is then trivial to take the transpose of a large matrix simply by changing the strides, with no need to re-order the matrix elements themselves.

This concept trivially extends to higher dimensions, as illustrated in Figure A1. Images can be stored with all intensities along the *x* axis stored one after the other, followed by the intensities for the next row along the *y* axis, followed by the intensities for the next slice along the *z* axis. For an image of size *n_x_*×*n_y_*×*n_z_*, this means the intensity at voxel (*x*, *y*, *z*) is stored at offset = *x* + *yn_x_*+ *zn_x_n_y_*; this corresponds to strides of [1 *n_x_n_x_n_y_*]. However, there is no particular reason for data to be stored in this order, and in fact different data strides are common. In many cases, the different image formats themselves assume different storage order conventions, which can cause confusion if not properly managed. Furthermore, the strides can also reflect the direction along which each axis is mapped, e.g., left-to-right versus right-to-left traversal, by allowing strides to be negative.

The extension to 4D can readily be appreciated in Figure A1, which also illustrates the case of volume-contiguous storage, i.e. when the intensities along the volume axis have adjacent offsets, with stride 1 for the volume index. Data stored in this way are sometimes referred to as *vector-valued* data: the data stored for each voxel is no longer a single intensity, but a *vector* of intensities. An example where this makes sense is when storing a colour image: it is then sensible to store the red, green & blue components of each pixel together. However, this is also advantageous in any situation where a vector of data is available for each voxel (e.g. all the fMRI or DWI intensities for a given voxel) and these data are processed independently per voxel (e.g. to perform a model fit). Computer hardware generally performs best when data are processed in the order they appear on file (or in memory): this avoids seek latency when reading from disk, allows burst transfers to and from the main system memory, leverages the CPU’s cache prefetching engine, and maximises usage of each line of the on-board CPU cache. Volume-contiguous strides therefore make sense for high-performance per-voxel operations on 4D data, which are common in diffusion MRI, functional MRI, perfusion MRI, T2 relaxometry, and many others domains.

To allow for all these different storage conventions and to enhance performance of the various analyses to be performed, *MRtrix3* supports arbitrary data strides for all supported formats, and for data held in RAM.

### 8.1 Symbolic strides

While the actual strides used to navigate the voxel information are unambiguous, they often involve large numbers, and depend on the exact image dimensions along each axis. To simplify the process of specifying and reporting strides, *MRtrix3* introduces the concepts of *symbolic* strides, and this is what is reported by tools such as *mrinfo*. With symbolic strides, the exact value of the stride is ignored, and all that matters is its magnitude relative to the other strides – i.e. its ranking (note that the sign of the strides is preserved). Hence, for the example shown in Figure A1, the strides would be reported as:

- *x → y → z → v*, actual strides = [1 3 9 18] ⇒ symbolic strides = [1 2 3 4]
- *y → x → z → v*, actual strides = [3 1 9 18] ⇒ symbolic strides = [2 1 3 4]
- *v → x → y → z*, actual strides = [2 6 18 1] ⇒ symbolic strides = [2 3 4 1]

Symbolic strides provide a simply means to specify the voxel ordering in a manner that is independent of the dimensions of the image. For example, if a 4D image had dimensions 96 × 128 × 64 × 32, with voxels stored in order *x → y → z → v* (as per the first example above), its actual strides would be [1 96 12288 786432]; yet its symbolic strides would be [1 2 3 4], matching the example above. If the user wishes to modify the strides of an image, it is therefore much more practical for them to specify them as symbolic strides, and let the software compute the actual strides given the dimensions of the image.

## 9 Appendix B: Example commands

The relatively simple examples below demonstrate stand-alone commands that compile / execute with *MRtrix3* without modification. They highlight a number of the powerful features available within *MRtrix3* that can be used with minimal coding effort, greatly expediting the development of image processing commands:

- Simple construction of an external *MRtrix3 module*, enabling compilation / execution of commands making use of *MRtrix3* functionalities without requiring any modification of *MRtrix3* itself;
- Consistent command-line interface, including checking validity of input parameters at the parsing stage;
- Compatibility with *any* image file format supported by the library;
- Simple construction of terminal feedback to user regarding command progress;
- Automated self-documentation that is consistent with all *MRtrix3* commands.

Two example commands are provided: One written in C++ (incorporating multi-threading), and one in Python. The functionality of these two demonstrative commands is identical:

- For any voxel where the value is less than a threshold (zero by default), set the intensity to zero; preserve the value of those voxels where the value is greater than the threshold.
- The threshold determining which image values should be clamped to zero may optionally be set manually by the user.

(Note that such a processing step can be achieved trivially using the *MRtrix3* command mrcalc; it is simply used here for demonstrative purposes)

### 9.1

#### 9.1.1 Example binary command

*Code* (file “demo_binary.cpp”):

This listing contains the C++ code to threshold an image, setting any intensities less than the specified threshold to zero.

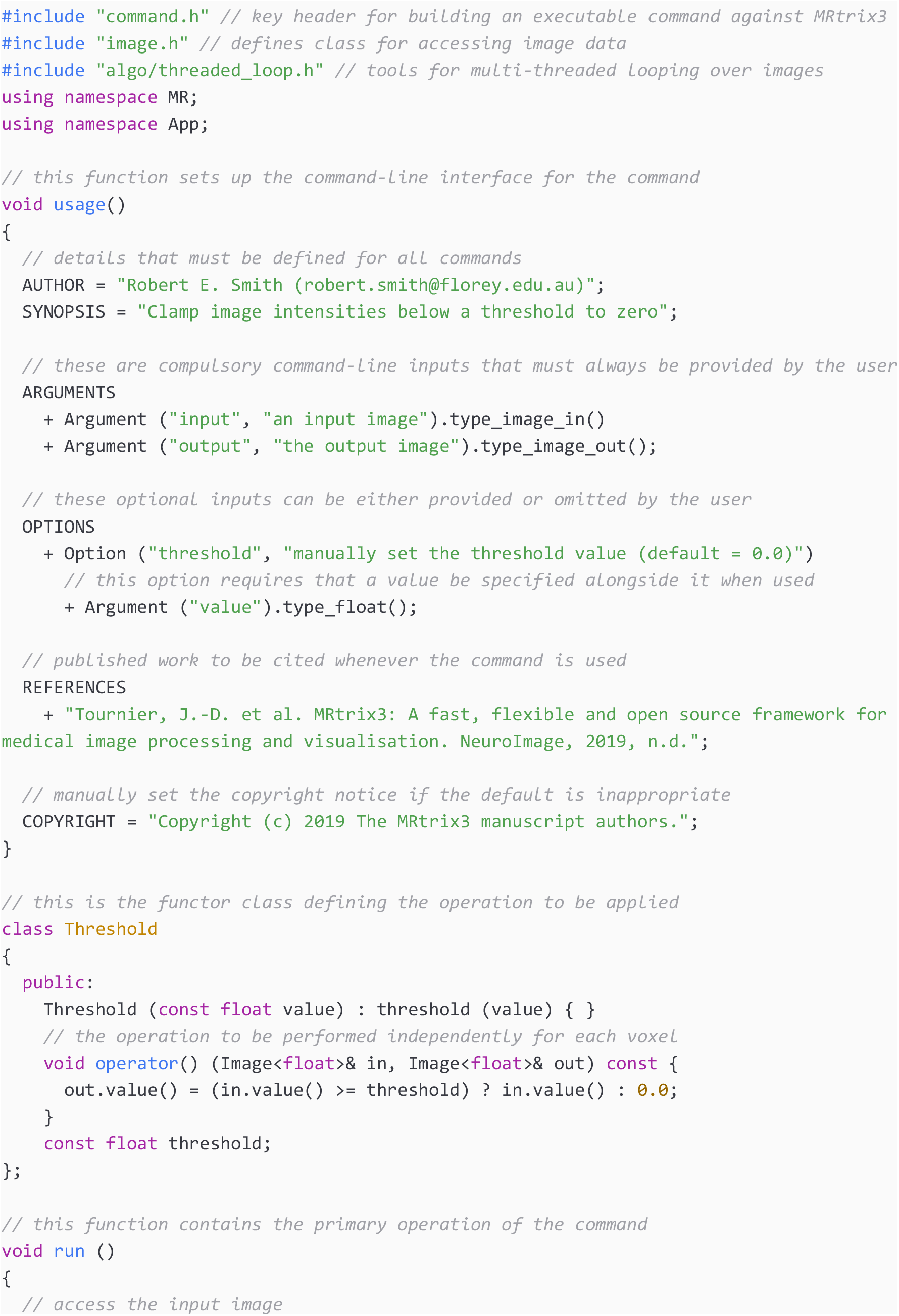
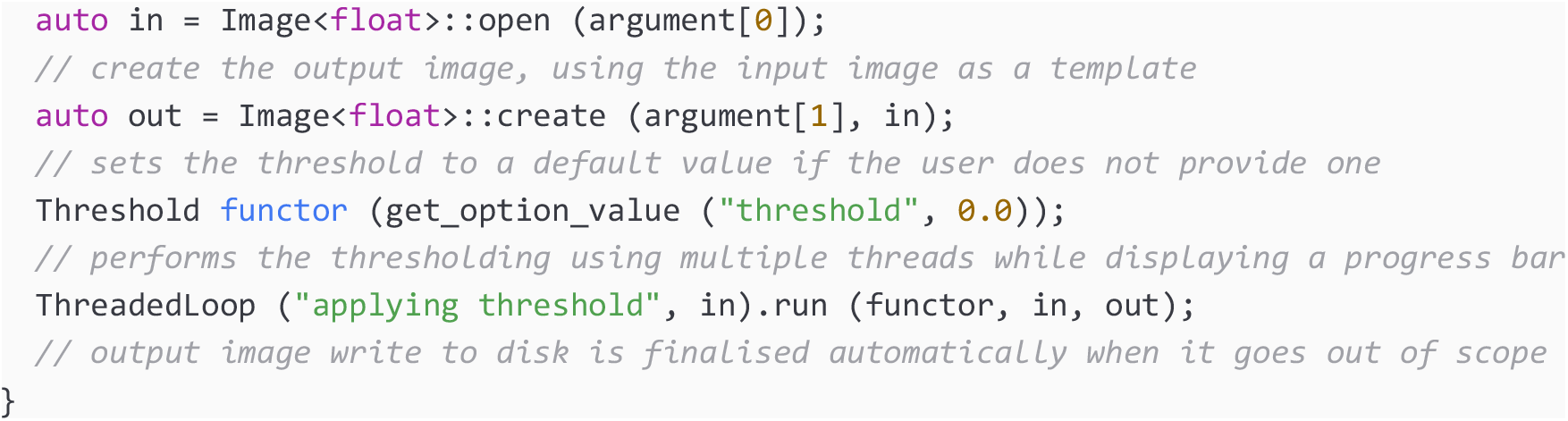

#### 9.1.2 Terminal usage

This listing shows the commands to invoke to build and execute the application for the code above. This assumes that the code sample has been saved to a file called ‘demo_binary.cpp’, currently residing in the user’s ‘Downloads’ folder.

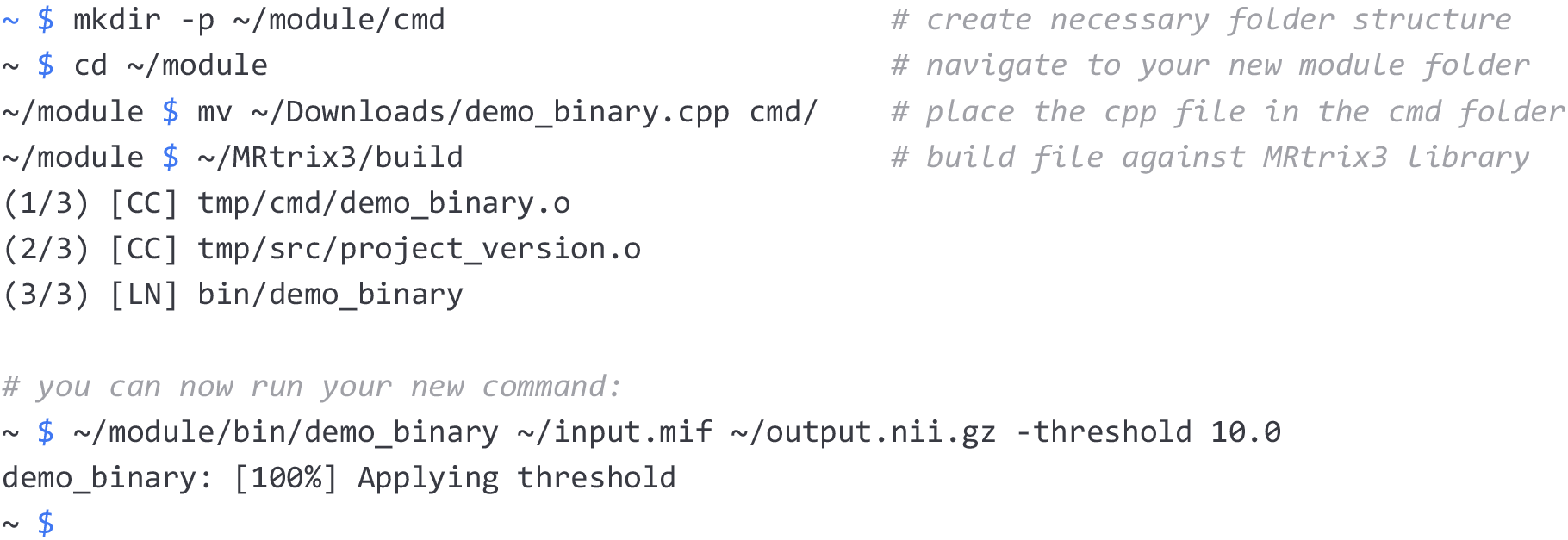

#### 9.1.3 Help page

This listing shows the help page print-out produced by invoking the ‘demo_binary’ command, compiled as above, with no arguments or with the ‘-help’ option. Note that the documentation matches the information provided by the developer in the ‘usage()’ function (see code listing in 9.1.1).

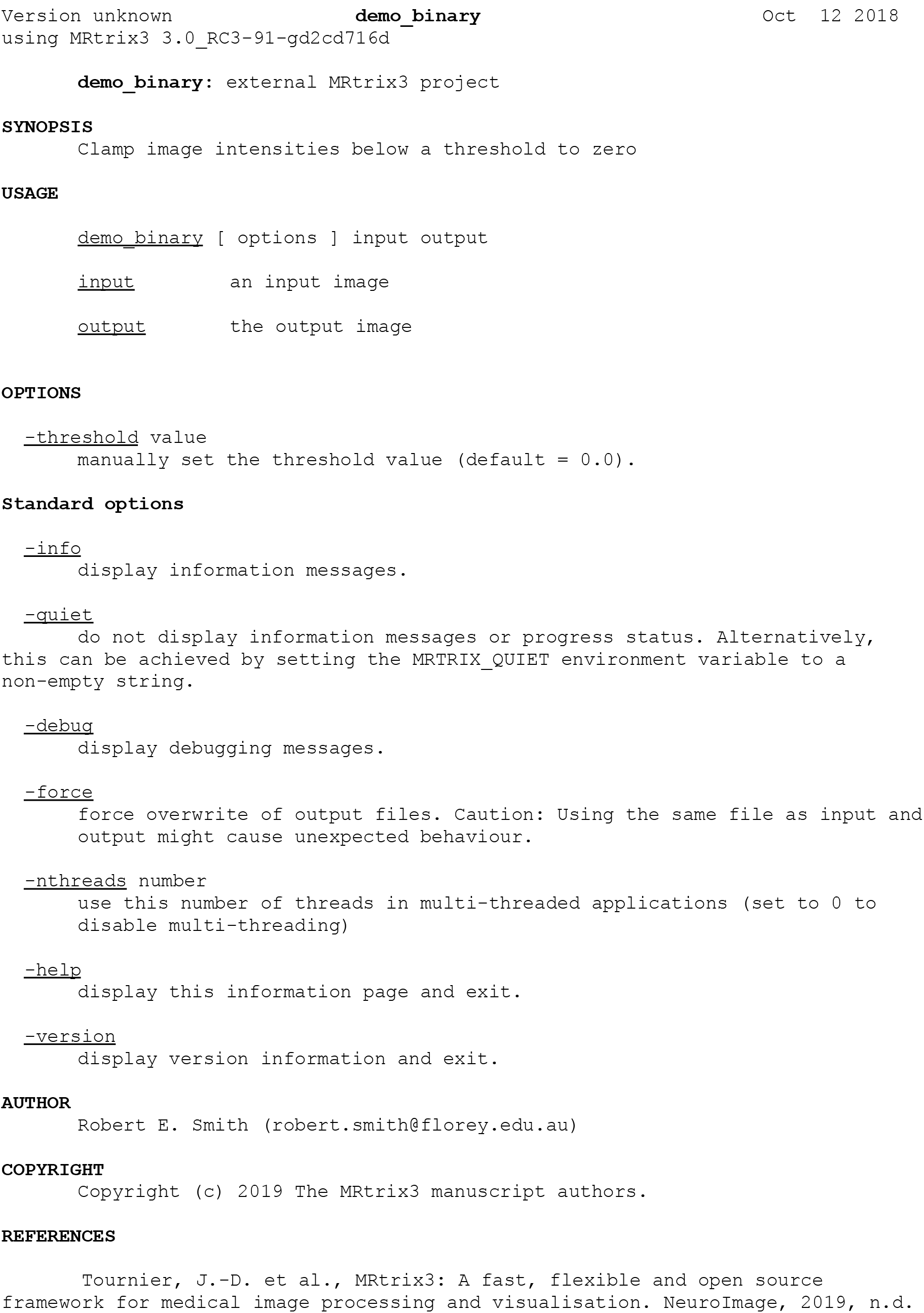

### 9.2 Example Python script

#### 9.2.1 Code (file “demo_python”)

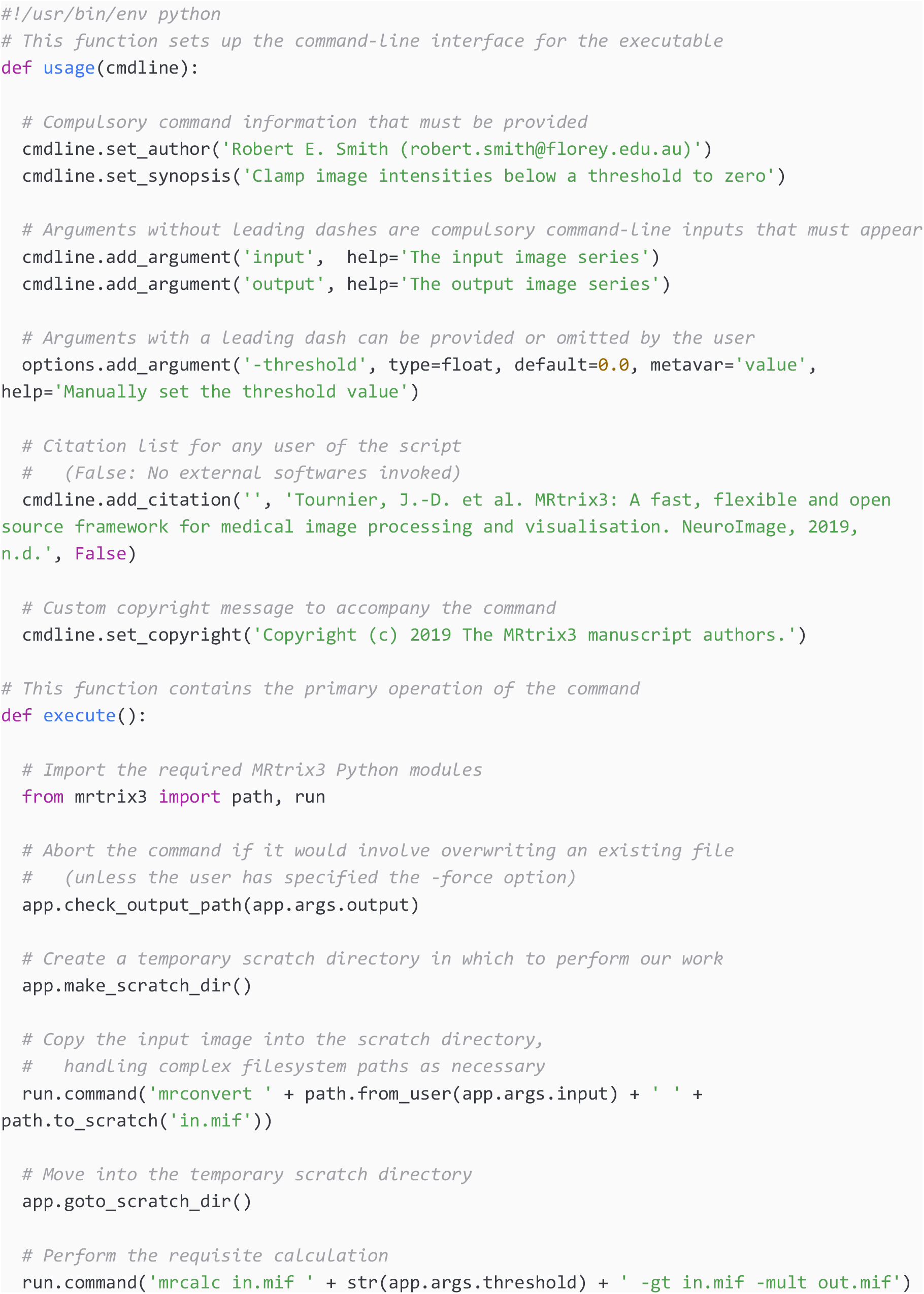
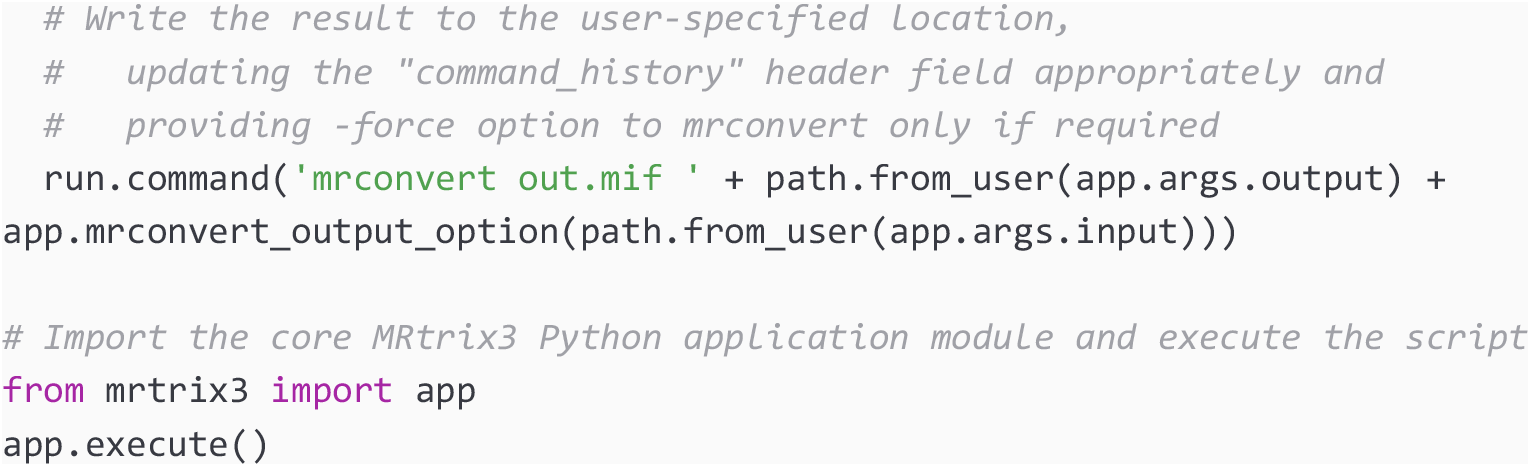

#### 9.2.2 Terminal usage

This listing shows the commands to invoke to set up and execute the application for the code above. This assumes that the code sample has been saved to a file called ‘demo_python’, currently residing in the user’s ‘Downloads’ folder.

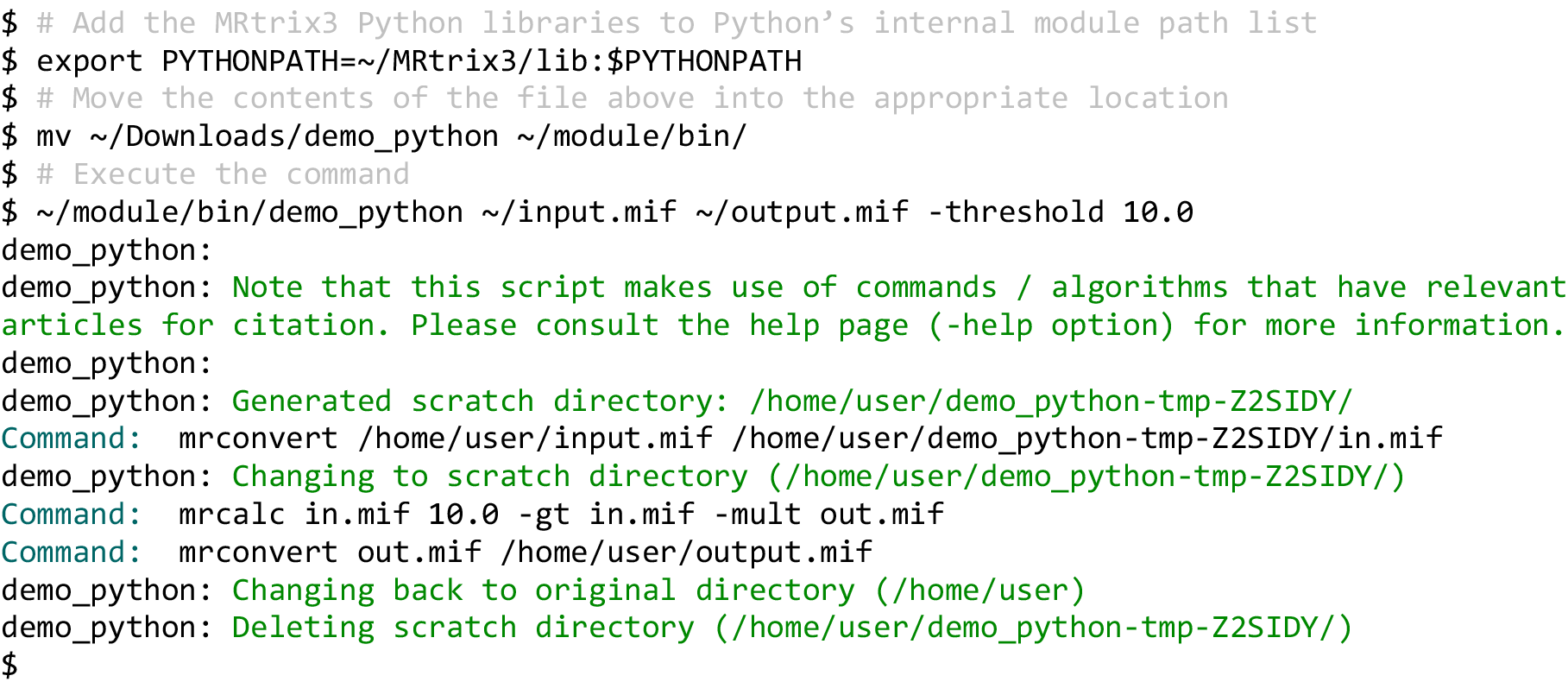

#### 9.2.3 Help page

The help page print-out produced by invoking the ‘demo_python’ command with no arguments or with the ‘-help’ option is practically identical to that produced by the ‘demo_binary’ command as shown in 9.1.3, with the exception being addition of the following:

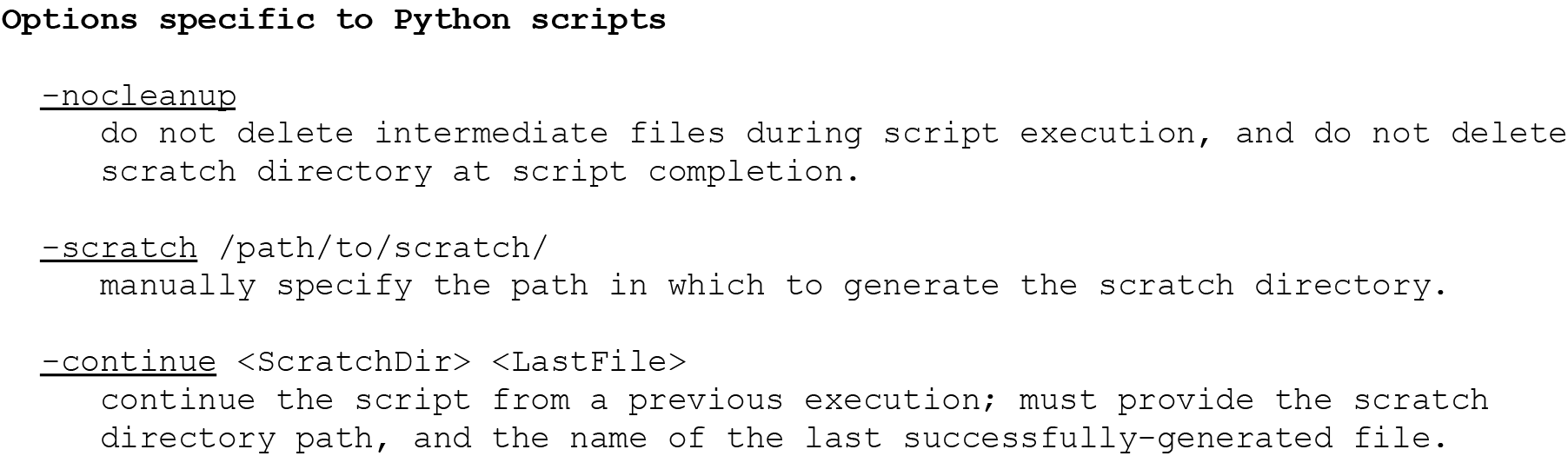

#### Footnotes

1 http://mozilla.org/MPL/2.0/

2 http://bids.neuroimaging.io

3 https://mrtrix.readthedocs.io

4 http://www.mrtrix.org

5 http://community.mrtrix.org

6 https://github.com/mrtrix3

7 http://www.open-std.org/jtc1/sc22/wg21/docs/standards

8 https://www.opengl.org/

9 https://www.python.org/

10 https://zlib.net/

11 http://eigen.tuxfamily.org/

12 https://www.qt.io/

13 https://www.msys2.org/

14 https://docs.microsoft.com/en-us/windows/wsl/about

15 https://www.dicomstandard.org/

16 https://rportal.mayo.edu/bir/ANALYZE75.pdf

17 https://nifti.nimh.nih.gov/nifti-1/

18 https://nifti.nimh.nih.gov/nifti-2/

19 https://surfer.nmr.mgh.harvard.edu/fswiki/FsTutorial/MghFormat

20 https://mrtrix.readthedocs.io/en/3.0_rc3/getting_started/image_data.html#mrtrix-image-formats-mih-mif

21 https://github.com/MRtrix3/mrtrix3/

22 https://github.com/MRtrix3/mrtrix3/issues

23 https://github.com/MRtrix3/test_data

24 https://travis-ci.org/MRtrix3/mrtrix3

25 https://ci.appveyor.com/project/MRtrix3/mrtrix3

26 http://www.mrtrix.org/developer-documentation/

27 http://www.doxygen.org/

28 http://www.mrtrix.org/developer-documentation/examples.html

29 https://www.opengl.org/

30 https://www.qt.io/

31 http://www.mrtrix.org/videos

32 https://mrtrix.readthedocs.io/en/3.0_rc3/concepts/fixels_dixels.html

33 https://mrtrix.readthedocs.io/en/3.0_rc3/getting_started/image_data.html#fixel-image-directory-format

34 http://www.libpng.org/pub/png/

